# rTMS to right parietal cortex decreases the precision of spatial priors in perceptual decision making

**DOI:** 10.1101/2023.08.23.554530

**Authors:** Arianna Zuanazzi, David Meijer, Uta Noppeney

## Abstract

Throughout life human observers make perceptual decisions under uncertainty guided by prior knowledge about the world’s causal structure and properties. According to Bayesian probability theory, optimal decision making relies on integrating prior knowledge with current sensory inputs, weighted by their relative precisions (i.e., inverse of uncertainty). Thus, representing uncertainty is essential for optimal decisions. Although extensive research suggests that human perceptual decision making follows Bayesian principles, the neural underpinnings of priors and their uncertainties remain elusive. In this five-day study, we employed psychophysics, Bayesian causal inference models, and offline low-frequency (i.e., 1Hz) transcranial magnetic stimulation (TMS) to probe the role of right intraparietal sulcus (IPS), a key region for spatial processing, in the integration of prior knowledge with auditory/visual inputs for spatial decisions. Model-free and Bayesian modelling analyses consistently showed a reduction in the precision of observers’ long-term spatial prior and in the influence of their previous spatial choices on their current spatial decisions for right IPS-TMS compared to sham-TMS. In contrast, the causal prior and the auditory/visual uncertainties remained unaffected. The results show that offline IPS-TMS can selectively reduce the precision or influence of observers’ long-term spatial prior and their short-term spatial expectations on perceptual decisions, without affecting their causal prior or sensory uncertainties (i.e., likelihood). Our findings provide causal evidence for the role of parietal cortex, situated at the top of the audiovisual spatial processing hierarchy, in encoding the uncertainty of spatial - but not causal - priors during perceptual decision-making.

**Significance statement:** Perceptual decisions under uncertainty are pervasive in everyday life. Substantial evidence suggests that humans perform perceptual decisions near-optimally. They combine sensory inputs with prior knowledge about the signals’ causal structure and locations weighted by their uncertainties. Yet, the neural underpinnings remain elusive. Combining psychophysics, Bayesian models, and offline low-frequency inhibitory transcranial magnetic stimulation (TMS), we provide causal evidence that the parietal cortex is crucial for encoding the uncertainty of spatial - but not causal - priors during perceptual decision-making. Intriguingly, observers relied less on their long-term and short-term prior spatial expectations after parietal-TMS, as compared to sham-TMS. These results provide important insights into the neural substrates of priors and their uncertainties as key ingredients for near-optimal decisions consistent with normative Bayesian principles.

## Introduction

In everyday life, we constantly need to make perceptual decisions under uncertainty by combining sensory inputs with prior knowledge about the world’s causal structure and properties (1–5). For instance, during a hike, we infer a rattlesnake’s location by integrating its rattling sound with the grass movement that it generates and our spatial expectations that rattlesnakes often hide in the tall grass near the path, as opposed to the busy path itself (i.e., spatial prior). Furthermore, our tendency to integrate signals across the senses is shaped by our prior expectation that they share a common cause. On windy days with continuous grass motion, we may be less inclined to integrate the rattling sound with nearby grass movement, compared to calm days when the grass motion likely originates from the rattlesnake (i.e., causal prior; (6)).

According to Bayesian probability theory, optimal decision making relies on integrating prior knowledge and new sensory inputs (i.e., likelihood), weighted by their relative precisions (i.e., inverse of uncertainty; (7)). Therefore, representing uncertainty is crucial for effective decision making. Growing research suggests that human perceptual decision making follows Bayesian principles. Most notably, human observers combine perceptual priors, such as prior beliefs about an object’s location or motion, with sensory inputs weighted by their relative precisions (8–12). Similarly, observers infer whether signals have common or independent sources by combining their prior knowledge about the signals’ causal structure with sensory correspondence cues, such as temporal synchrony or spatial colocation (6, 13–15). According to Bayesian Causal inference models, this causal prior also shapes observers’ perceptual (e.g., spatial) estimates, by modulating the balance between sensory integration and segregation. For instance, a strong common source causal prior increases the dominance of the perceptual estimate that integrates audiovisual signals on observers’ percepts (6, 16).

Neuroimaging research has shown that the brain accomplishes Bayesian causal inference by dynamically computing multiple perceptual estimates along the cortical hierarchy (16–22). In the domain of spatial processing, early sensory cortices form spatial estimates that segregate the auditory and visual signals; posterior parietal cortices then merge auditory and visual signals weighted by their relative precisions; at the top of the hierarchy, anterior parietal cortices form spatial estimates that flexibly combine audiovisual inputs in accordance with the principles of Bayesian causal inference: they integrate audiovisual signals that are spatially proximate and thus likely share a common cause, but segregate them otherwise; in contrast, dorsolateral prefrontal cortices determine whether signals originate from common or independent sources (23). Thus, parietal cortices appear to compute spatial representations that are informed by causal inference decisions in dorsolateral prefrontal cortices.

Despite this abundant behavioral and neuroimaging evidence that human observers weigh sensory evidence and prior knowledge by their precisions, the neural underpinnings of priors and their uncertainties remain elusive. Leading theoretical accounts propose that priors and their uncertainties are represented within sensory pathways (24, 25), whereas some previous neuroimaging work pointed to neural circuitries outside these pathways (7). Importantly, neuroimaging as a correlational method cannot ascertain whether a region plays a causal role or is merely implicitly activated.

In this five-day study, we employed psychophysics, Bayesian causal inference models, and offline low-frequency (i.e., 1Hz) transcranial magnetic stimulation (TMS) to probe the causal role of anterior right intraparietal sulcus (IPS) in the integration of causal and spatial prior knowledge with spatially congruent or disparate audiovisual and unisensory auditory/visual inputs for spatial decisions. This multisensory setting enabled us to investigate whether right IPS-TMS (vs. sham-TMS) perturbs the integration of sensory evidence (i.e., likelihood) with spatial or causal priors (i.e., common vs. separate causes) formed at long- and short timescales. Based on previous neuroimaging work, we expected IPS-TMS (vs. sham-TMS) to selectively reduce the influence of spatial - but not causal - priors on observers’ perceptual decisions (16, 23).

## Results

Twenty-two participants were presented with synchronous visual and auditory signals sampled independently from four locations (i.e., -8°, -16°, -24°, -32°) in their left hemifield. This design resulted in four levels of audiovisual spatial disparity: 0°, 8°, 16° or 24° (**Fig.1A** and **1B**). Additionally, unisensory auditory and visual stimuli were presented independently from the same four locations. Participants were required to localize either the visual or the auditory stimulus component under central fixation. Before the task, offline low-frequency (i.e., 1Hz) repetitive TMS was applied to their right anterior intraparietal sulcus. The coil was aligned either parallel (i.e., IPS-TMS) or perpendicular (i.e., sham-TMS) to participants’ scalp on separate days (**Fig.1C** and **Materials and Methods**).

**Figure 1.**
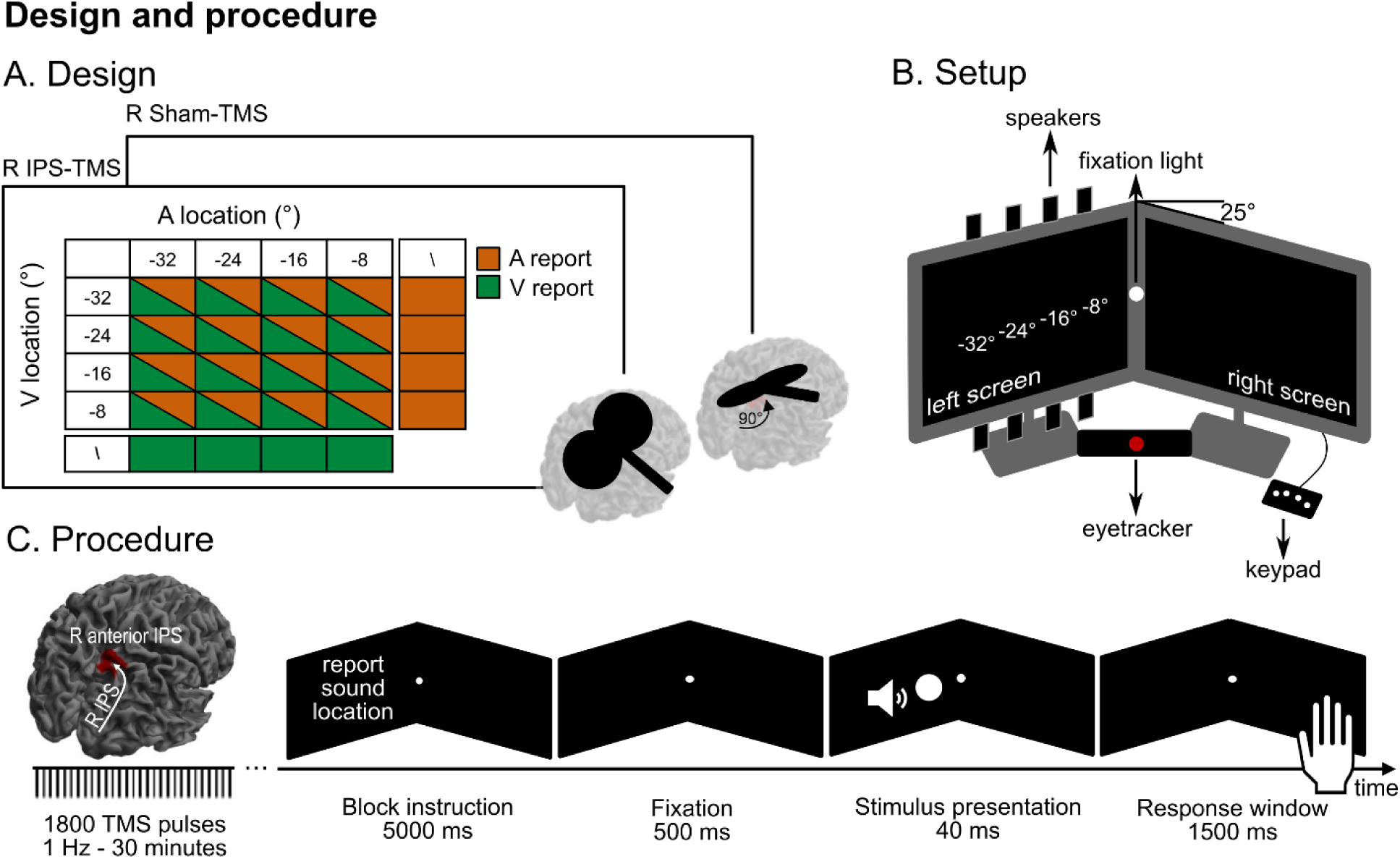
Experiment design, setup and procedure. **A.** The 4 × 4 × 2 × 2 factorial design manipulated (i) location of the auditory stimulus (-8°, -16°, - 24°, -32°), (ii) location of the visual stimulus (-8°, -16°, -24°, -32°), (iii) relevant sensory modality (report auditory or visual location, orange and green filling) across blocks, and (iv) TMS (IPS-TMS or sham-TMS) manipulated across sessions (on different days). Each relevant sensory modality block also included auditory or visual unisensory trials (‘\’ in the figure). Sham-TMS was performed with the coil tilted by 90° with the wings touching the scalp over right anterior IPS (**Materials and Methods**). **B.** Experimental setup: two LCD monitors tilted by -25° (monitor on the left) and +25° (monitor on the right) perpendicular to participants’ eye gaze direction. A fixation light was mounted in the middle of the two monitors at 0° elevation. Auditory and visual stimuli were presented at -8°, -16°, -24° and -32° along the azimuth. To ensure fixation, eye-movements were recorded via an EyeLink eyetracking system, throughout the duration of the experiment. **C.** The task was preceded by 30 minutes of offline TMS stimulation at 1 Hz. During the task, participants were presented with blocks of audiovisual signals and unisensory auditory or visual signals, originating from the four possible locations. Before each block, a cue indicated the relevant sensory modality (i.e., whether to report the location of the sound or the flash). The flash consisted of a white circle (radius: 2.5° visual angle) and the sound was a burst of white noise presented through two of eight loudspeakers that were attached to the top and bottom of the left screen. Stimuli duration was 40 ms for both sounds and flashes. Immediately after a response (pressing one of four buttons), or if no response was given within 1500 ms, the next trial would start with 500 ms pre-stimulus period at which participants fixated the central light.

Overall, observers localized auditory and visual signals reliably as indicated by significant correlation coefficients between participants’ responses and the true stimulus locations on unisensory and audiovisual congruent trials. On average, participants’ accuracy was ρ = 0.84 for auditory report and ρ = 0.95 for visual report. A 2 (*TMS*: sham-TMS vs. IPS-TMS) × 2 (*Relevant sensory modality*: report Auditory vs. Visual stimuli) × 2 (*Type*: Auditory/Visual vs. Audiovisual congruent) repeated measures ANOVA revealed a significant main effect of relevant sensory modality (F(1,21) = 108.39, *p* < 0.001, η_p_^2^ = 0.84) but no significant interactions, with higher correlation coefficients for report Visual than report Auditory stimuli.

The main focus of our study was to determine whether and how TMS to anterior parietal cortices influences the spatial perception of unisensory auditory/visual and audiovisual stimuli. We assessed this by comparing observers’ spatial localization performance under right IPS-TMS and sham-TMS, using model-free indices (‘model-free analysis’) and parameters of a Bayesian Causal Inference model fitted to participants’ responses (‘model-based analysis’).

### ‘Model-free’ (MF) analyses

#### No effect of IPS-TMS on σ_MF_S_ as an index for sensory noise

We initially examined whether IPS-TMS, compared to sham-TMS, affects sensory noise – either auditory or visual – in the unisensory or the audiovisual congruent conditions. To estimate sensory noise, we computed the spread (standard deviation, σ_MF_s_) of the reported sound or flash locations for each stimulus location and then pooled these values over locations. This standard deviation served as a model-free index for sensory noise, which corresponds to the sensory noise variance estimated in the Bayesian Causal inference model (i.e., σ_S,_ see **Materials and Methods**). A 2 (*TMS*: sham-TMS vs. IPS-TMS) × 2 (*Relevant sensory modality*: report Auditory vs. Visual stimuli) × 2 (*Type*: Auditory/Visual vs. Audiovisual congruent) repeated measures ANOVA on the sensory noise σ_MF_S_ revealed a significant main effect of relevant sensory modality (**Table 1A** and **Fig.2A**). As expected, the spatially more reliable visual stimuli were associated with less sensory noise than the auditory stimuli. We also observed a small non-significant effect of stimulus type, indicating a trend towards smaller auditory response variance on audiovisual stimuli with auditory report than unisensory auditory stimuli (**Table 1A** and **Fig.2A**). This noise reduction is expected for congruent trials due to audiovisual integration. Importantly, however, low-frequency IPS-TMS had no significant effect on observers’ sensory noise. For a comprehensive list of results see **Table 1**.

**Figure 2.**
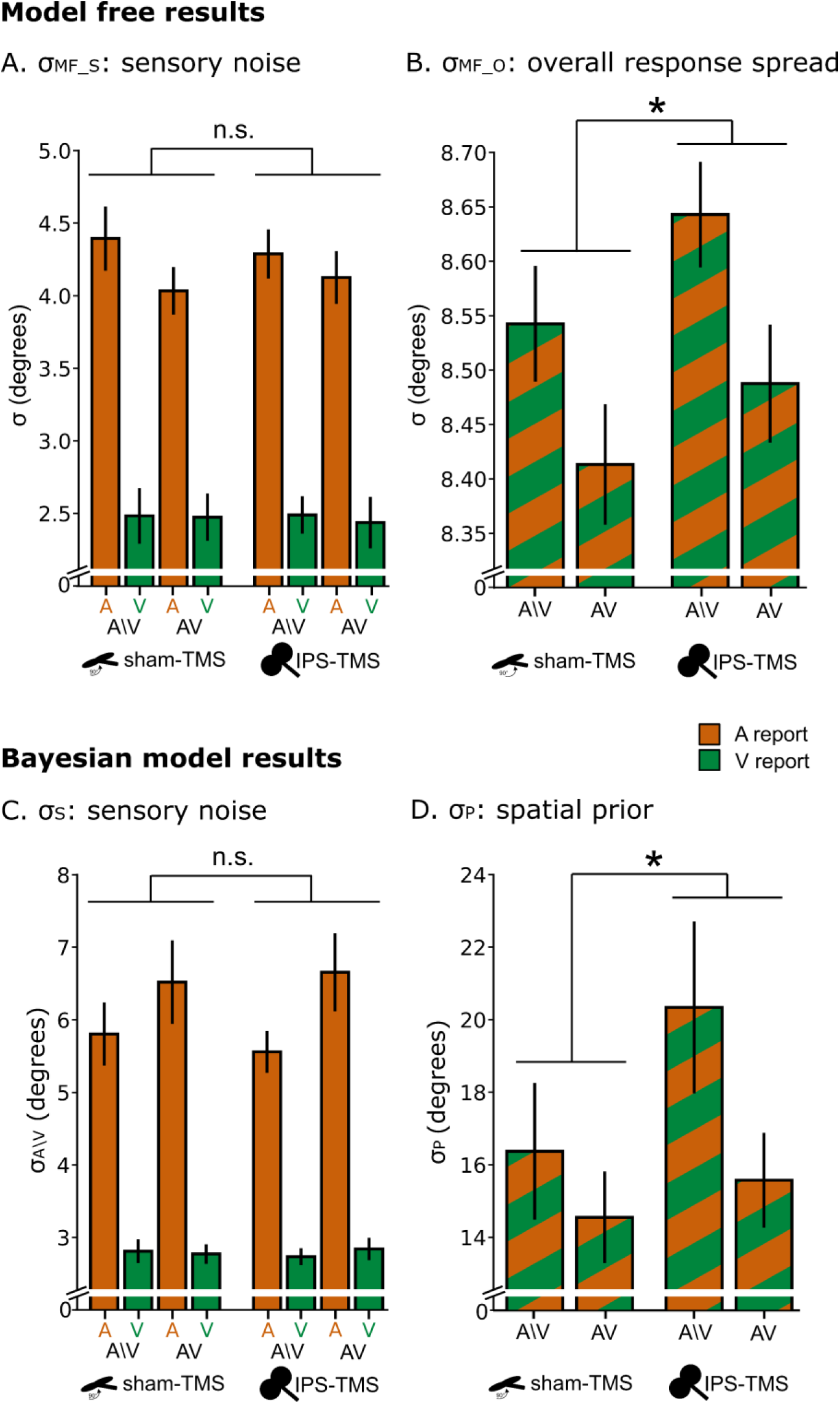
Results of the model-free (MF) analysis and of the Bayesian Causal Inference (BCI) modelling analysis. Model-free results: **A.** Across participants’ mean (±SEM) of sensory noise standard deviations. **B.** Across participants’ mean (±SEM) of overall response spread or standard deviation. Bayesian modelling results: **C.** Across participants’ mean (±SEM) of sensory noise standard deviation. **D.** Across participants’ mean (±SEM) of the spatial prior standard deviation. Only significant results for TMS are indicated with asterisks. Please refer to **Table 1A-C** for a comprehensive list of significant effects. ** *p* < 0.01, * *p* < 0.05.

**Table 1.**
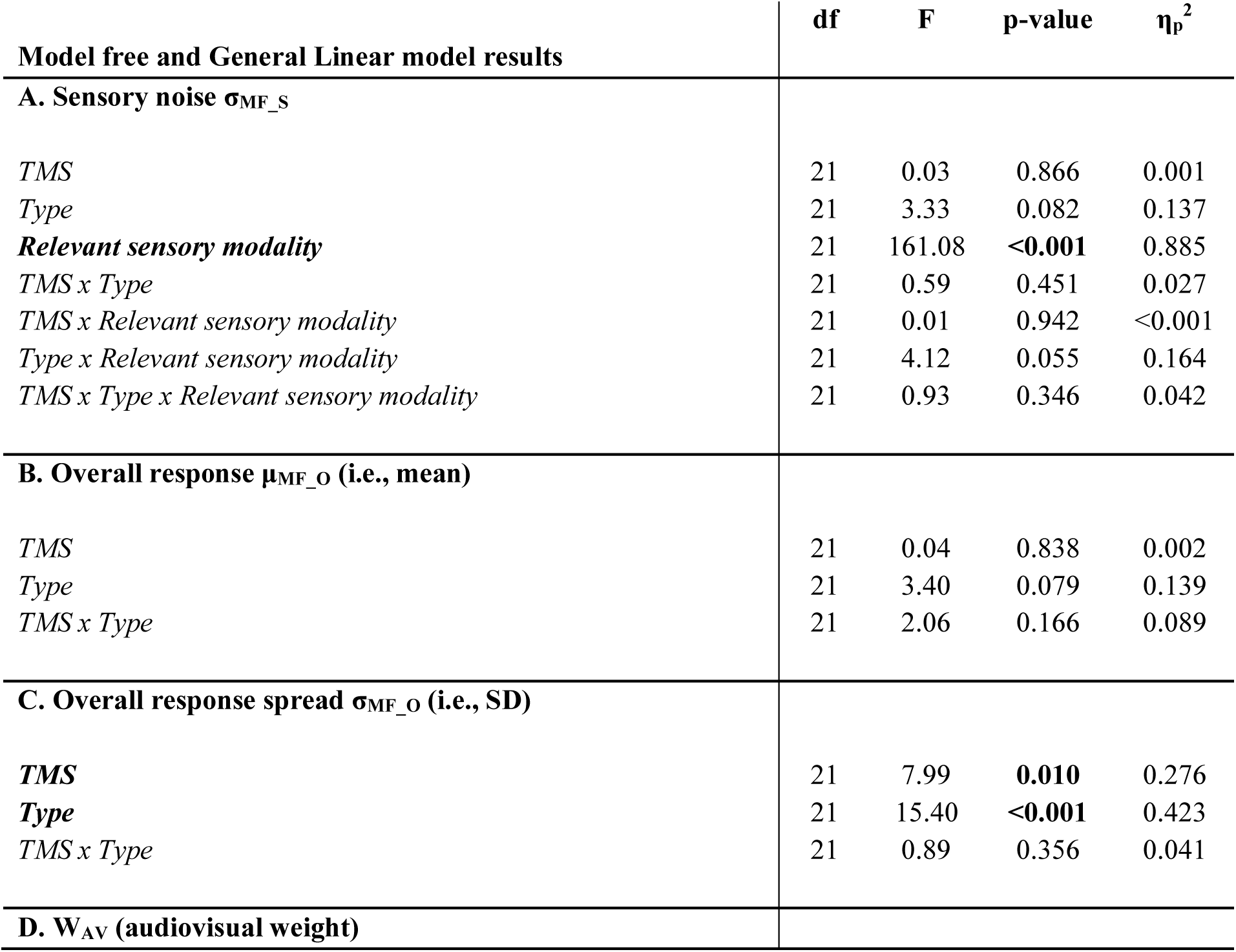
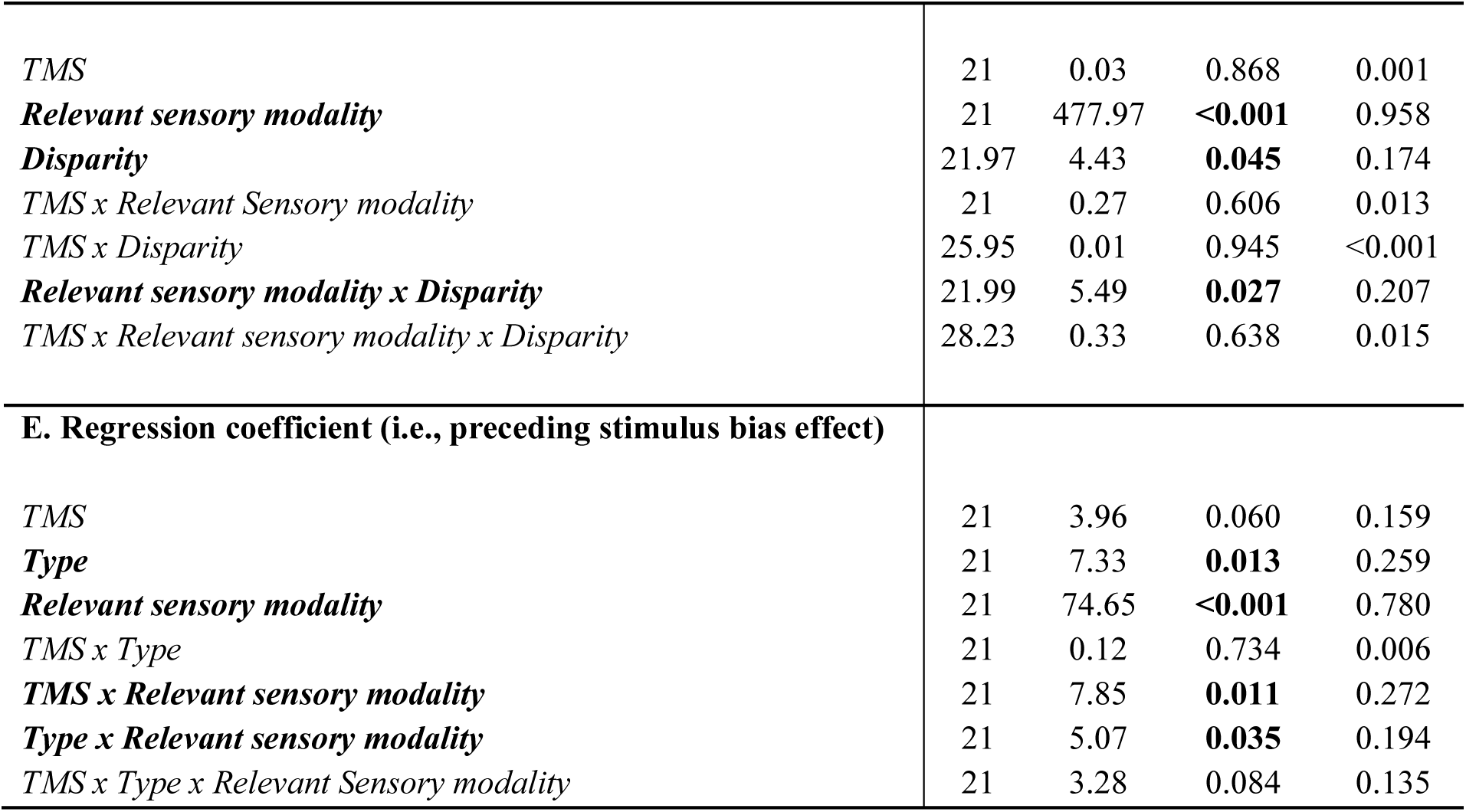
Summary of results of ‘Model-Free’ (MF) analyses. *TMS*: sham-TMS vs. IPS-TMS; *Relevant sensory modality*: report Auditory vs. Visual stimuli; *Type*: Auditory/Visual vs. Audiovisual congruent; *Disparity*: 8°, 16°, 24°. Significant factors and their *p*-values are highlighted in bold. Sphericity correction: Greenhouse-Geisser.

| | df | F | p-value | $\eta_p^2$ |
| --- | --- | --- | --- | --- |
| **Model free and General Linear model results** |  |  |  |  |
| **A. Sensory noise $\sigma_{MF\_S}$** |  |  |  |  |
| *TMS* | 21 | 0.03 | 0.866 | 0.001 |
| *Type* | 21 | 3.33 | 0.082 | 0.137 |
| ***Relevant sensory modality*** | 21 | 161.08 | **<0.001** | 0.885 |
| *TMS x Type* | 21 | 0.59 | 0.451 | 0.027 |
| *TMS x Relevant sensory modality* | 21 | 0.01 | 0.942 | <0.001 |
| *Type x Relevant sensory modality* | 21 | 4.12 | 0.055 | 0.164 |
| *TMS x Type x Relevant sensory modality* | 21 | 0.93 | 0.346 | 0.042 |
| **B. Overall response $\mu_{MF\_O}$ (i.e., mean)** |  |  |  |  |
| *TMS* | 21 | 0.04 | 0.838 | 0.002 |
| *Type* | 21 | 3.40 | 0.079 | 0.139 |
| *TMS x Type* | 21 | 2.06 | 0.166 | 0.089 |
| **C. Overall response spread $\sigma_{MF\_O}$ (i.e., SD)** |  |  |  |  |
| ***TMS*** | 21 | 7.99 | **0.010** | 0.276 |
| ***Type*** | 21 | 15.40 | **<0.001** | 0.423 |
| *TMS x Type* | 21 | 0.89 | 0.356 | 0.041 |
| **D. $W_{AV}$ (audiovisual weight)** |  |  |  |  |
| *TMS* | 21 | 0.03 | 0.868 | 0.001 |
| ***Relevant sensory modality*** | 21 | 477.97 | **<0.001** | 0.958 |
| ***Disparity*** | 21.97 | 4.43 | **0.045** | 0.174 |
| *TMS x Relevant Sensory modality* | 21 | 0.27 | 0.606 | 0.013 |
| *TMS x Disparity* | 25.95 | 0.01 | 0.945 | <0.001 |
| ***Relevant sensory modality x Disparity*** | 21.99 | 5.49 | **0.027** | 0.207 |
| *TMS x Relevant sensory modality x Disparity* | 28.23 | 0.33 | 0.638 | 0.015 |
| **E. Regression coefficient (i.e., preceding stimulus bias effect)** |  |  |  |  |
| *TMS* | 21 | 3.96 | 0.060 | 0.159 |
| ***Type*** | 21 | 7.33 | **0.013** | 0.259 |
| ***Relevant sensory modality*** | 21 | 74.65 | **<0.001** | 0.780 |
| *TMS x Type* | 21 | 0.12 | 0.734 | 0.006 |
| ***TMS x Relevant sensory modality*** | 21 | 7.85 | **0.011** | 0.272 |
| ***Type x Relevant sensory modality*** | 21 | 5.07 | **0.035** | 0.194 |
| *TMS x Type x Relevant Sensory modality* | 21 | 3.28 | 0.084 | 0.135 |

#### Effect of IPS-TMS on the observers’ overall response spread, but not mean

Next, we investigated whether IPS-TMS shifts the mean (i.e., µ) or changes the spread (i.e., standard deviation, σ) of observers’ overall response distribution. A shift in the mean of observers’ overall response distribution could suggest that TMS introduces a spatially lateralized bias, while a change in the overall spread of the response distribution may result from changes in spatial prior and/or sensory noise variance. A 2 (*TMS*: sham-TMS vs. IPS-TMS) × 2 (*Type*: Auditory/Visual vs. Audiovisual) repeated measures ANOVA on observers’ overall response mean (i.e., µ_MF_O_) did not indicate any significant effects of TMS or stimulus type (**Table 1B**). However, the same analysis on the response distribution’s spread (i.e., σ_MF_O_) demonstrated significant effects of stimulus type and TMS (**Table 1C** and **Fig.2B**). Observers’ response distributions under sham-TMS were more focal and centered on the distribution’s mean compared to those under IPS-TMS. Given that TMS did not affect sensory noise (**Table 1A** and **Fig.2A)**, this broadening of the response distribution after IPS-TMS provides initial evidence for a decrease in the precision of observers’ central prior and hence its reduced influence (or weight) during spatial perception, across both unisensory auditory/visual and audiovisual conditions.

We also observed a wider spread of the response distribution for unisensory auditory/visual as compared to audiovisual blocks (i.e., main effect of stimulus type; **Table 1C** and **Fig.2B**). This might indicate a greater reliance on spatial priors under audiovisual stimulation, potentially due to higher attentional demands in those audiovisual blocks that present spatially congruent and incongruent signals in a randomized fashion. For a comprehensive list of results see **Table 1**.

To ensure that these effects on observers’ response distribution were not driven by crossmodal biases, we repeated the analysis constrained to unisensory and congruent audiovisual trials only. Importantly, this more constrained analysis replicated the significant effects of TMS (F(1,21) = 5.81, *p* = 0.025, η_p_^2^ = 0.22) and stimulus type (F(1,21) = 6.79, *p* = 0.016, η_p_^2^ = 0.24).

#### No effects of IPS-TMS on audiovisual weight (W_AV_)

To assess how observers weigh and integrate audiovisual signals into spatial percepts, we computed the audiovisual weight index W_AV_ separately for auditory and visual reports, across the three levels of spatial disparity (i.e., 8°, 16°, 24°). The audiovisual weight index ranges from one (indicating pure visual influence) to zero (indexing pure auditory influence; see **Materials and Methods**). As expected under Bayesian causal inference models (6, 15), observers did not fuse auditory and visual signals into one unified percept. Instead, they gave a stronger weight to the auditory signal when asked to report the sound location and to the visual signal when asked to report the flash location. A 2 (*TMS*: sham-TMS vs. IPS-TMS) × 2 (*Relevant sensory modality*: Auditory vs. Visual report) × 3 (*Disparity*: 8°, 16°, 24°) repeated measures ANOVA on the audiovisual weight index W_AV_ confirmed a significant main effect of relevant sensory modality (**Table 1D** and **Fig.3A**). Moreover, and in line with Bayesian causal inference and previous research (6, 15, 19, 26), this difference in W_AV_ between auditory and visual report increased with spatial disparity (i.e., significant interaction between relevant sensory modality and disparity; **Table 1D** and **Fig.3A**). At large spatial disparities, where it is less likely for audiovisual signals to originate from common sources, audiovisual interactions diminished and wAV was closer to zero for auditory reports. Nevertheless, even for the largest spatial disparity (24°) Student t tests confirmed that wAV was larger than zero for auditory reports (t(21) = 2.34, *p* = 0.029 and t(21) = 2.22, *p* = 0.038 for Sham-TMS and IPS-TMS, respectively; with wAV ≈ 6% for both).

**Figure 3.**
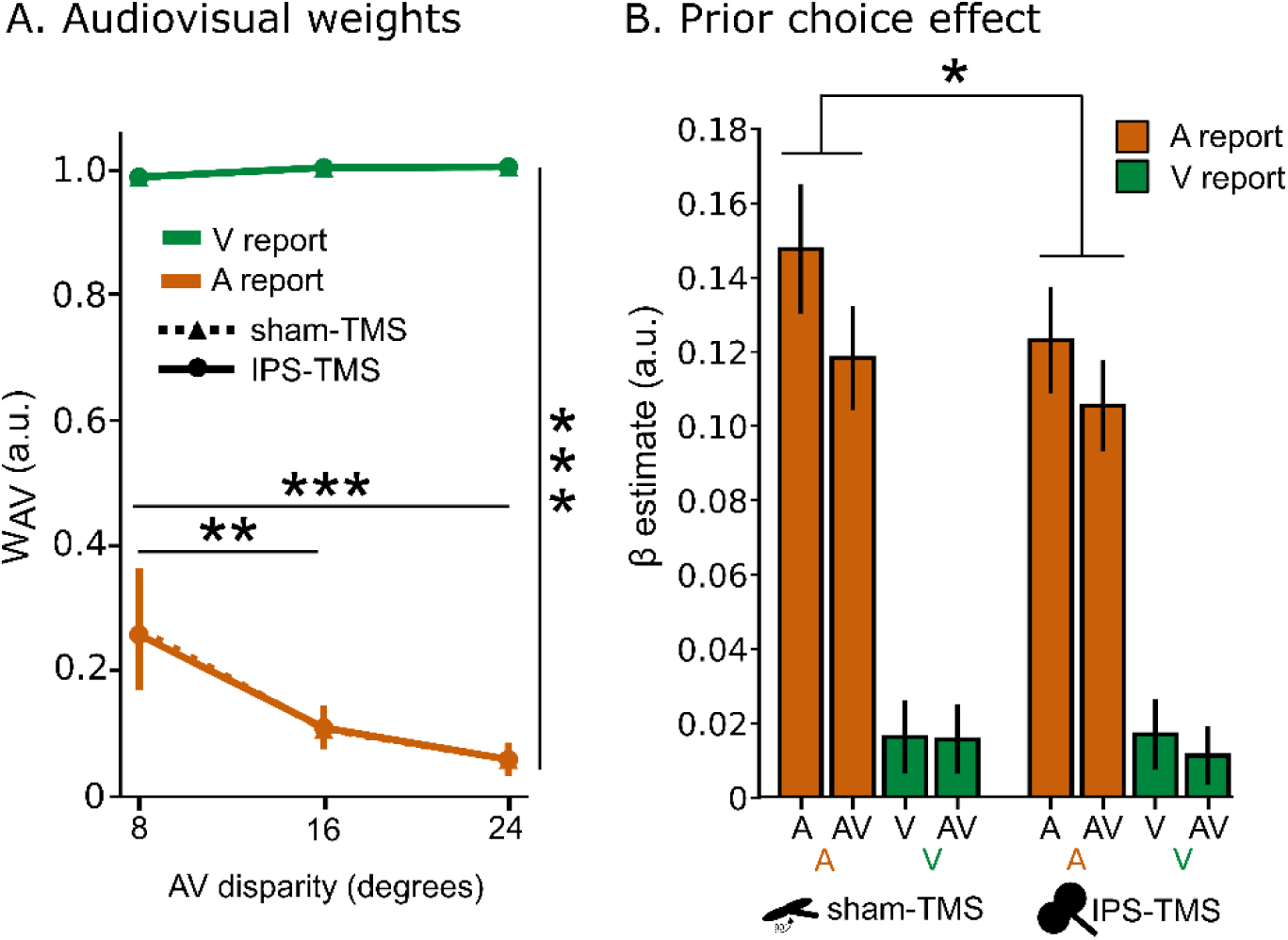
Audiovisual bias and spatial bias from previous response. **A.** Across participants’ mean (±SEM) audiovisual weights for auditory and visual reports, for each audiovisual disparity. W_AV_ = 1 indicates that the observer’s spatial report relies exclusively on the location of the visual signal. W_AV_ = 0 indicates that the observer’s spatial report relies exclusively on the location of the auditory signal. **B.** Regression coefficients defining the bias of the previous stimulus. Please refer to **Table 1D-E** for a comprehensive list of significant effects. *** *p* < 0.001, ** *p* < 0.01, * *p* < 0.05.

According to precision weighted integration, we would expect the perceived sound location to be shifted more towards the true visual location than the perceived visual location towards the true auditory location. In line with this conjecture, the audiovisual weight index W_AV_ was significantly different from zero under auditory report (see above), but not significantly different from 1 under visual report (*p* > 0.05 for all six conditions).

Critically, however, IPS-TMS had no significant effect on the audiovisual weight index (i.e., no significant main effect or interaction, **Table 1D**), suggesting that IPS-TMS does not influence how the brain weighs and combines audiovisual spatial signals. For a comprehensive list of results see **Table 1**.

In summary, our model-free analysis revealed that, compared to sham-TMS, IPS-TMS influences perceptual inference by enhancing observers’ central response tendencies with no effect on their sensory noise or audiovisual interactions. These findings point towards a possible impact of IPS-TMS on the precision (or variance) of observers’ spatial prior. Because model-free analyses do not allow for a complete dissociation of prior and sensory evidence, we next turn to model-based analyses and fit the Bayesian causal inference model to observers’ response distributions.

### Model-based analysis: Bayesian causal inference (BCI) model

We fitted Bayesian Causal Inference models (BCI) to audiovisual response distributions for each participant in both sham-TMS and IPS-TMS conditions. This entailed estimating five parameters: sensory noise (σ_A_ and σ_V_), the mean and standard deviations of the spatial prior (μ_P_ and σ_P_) and the causal prior (p_common_) that quantifies observers’ prior tendency to bind audiovisual signals (**Table 2** and see **Materials and Methods**). For the unisensory auditory and visual conditions, we fitted the first four parameters only, while fixing the causal prior to zero which indicates full segregation. Similar to our model-free analysis, we entered these parameter estimates into repeated measures ANOVAs at the group level, to assess effects of TMS, relevant sensory modality, and stimulus type.

**Table 2.** Summary of parameters estimates of the Bayesian Causal Inference model. Group mean (SEM).

| Bayesian model parameter estimates | Unisensory |  | Audiovisual |  |
| --- | --- | --- | --- | --- |
|  | Sham-TMS | IPS-TMS | Sham-TMS | IPS-TMS |
| **Auditory sensory noise ($\sigma_A$)** | 5.80 (0.44) | 5.56 (0.29) | 6.52 (0.57) | 6.66 (0.54) |
| **Visual sensory noise ($\sigma_V$)** | 2.81 (0.16) | 2.73 (0.12) | 2.77 (0.13) | 2.84 (0.15) |
| **Spatial prior mean ($\mu_P$)** | -23.9 (3.13) | -28.2 (6.61) | -22.8 (3.93) | -24.8 (3.17) |
| **Spatial prior standard deviation ($\sigma_P$)** | 16.4 (1.89) | 20.3 (2.37) | 14.5 (1.26) | 15.6 (1.31) |
| **Causal prior ($p_{\text{common}}$)** | N/A | N/A | 0.11 (0.03) | 0.13 (0.04) |

#### IPS-TMS does not influence sensory noise standard deviations (σ_A_ and σ_V_)

A 2 (*TMS*: sham-TMS vs. IPS-TMS) × 2 (*Relevant sensory modality*: report Auditory vs. Visual stimuli) × 2 (*Type*: Auditory/Visual vs. Audiovisual) repeated measures ANOVA revealed a significant main effect of relevant sensory modality and stimulus type as well as a significant interaction between stimulus type and relevant sensory modality (**Table 3A** and **Fig.2C**). As expected, and consistent with our model-free results, the visual noise was smaller than the auditory noise. Moreover, the auditory noise was smaller for auditory stimuli than audiovisual stimuli, likely due to the increased attentional demands resulting from the random presentation of spatially congruent and incongruent signals in the audiovisual blocks. Crucially, IPS-TMS did not influence the auditory or the visual noise (**Table 3A** and **Fig.2C**), corroborating our model-free results.

**Table 3.**
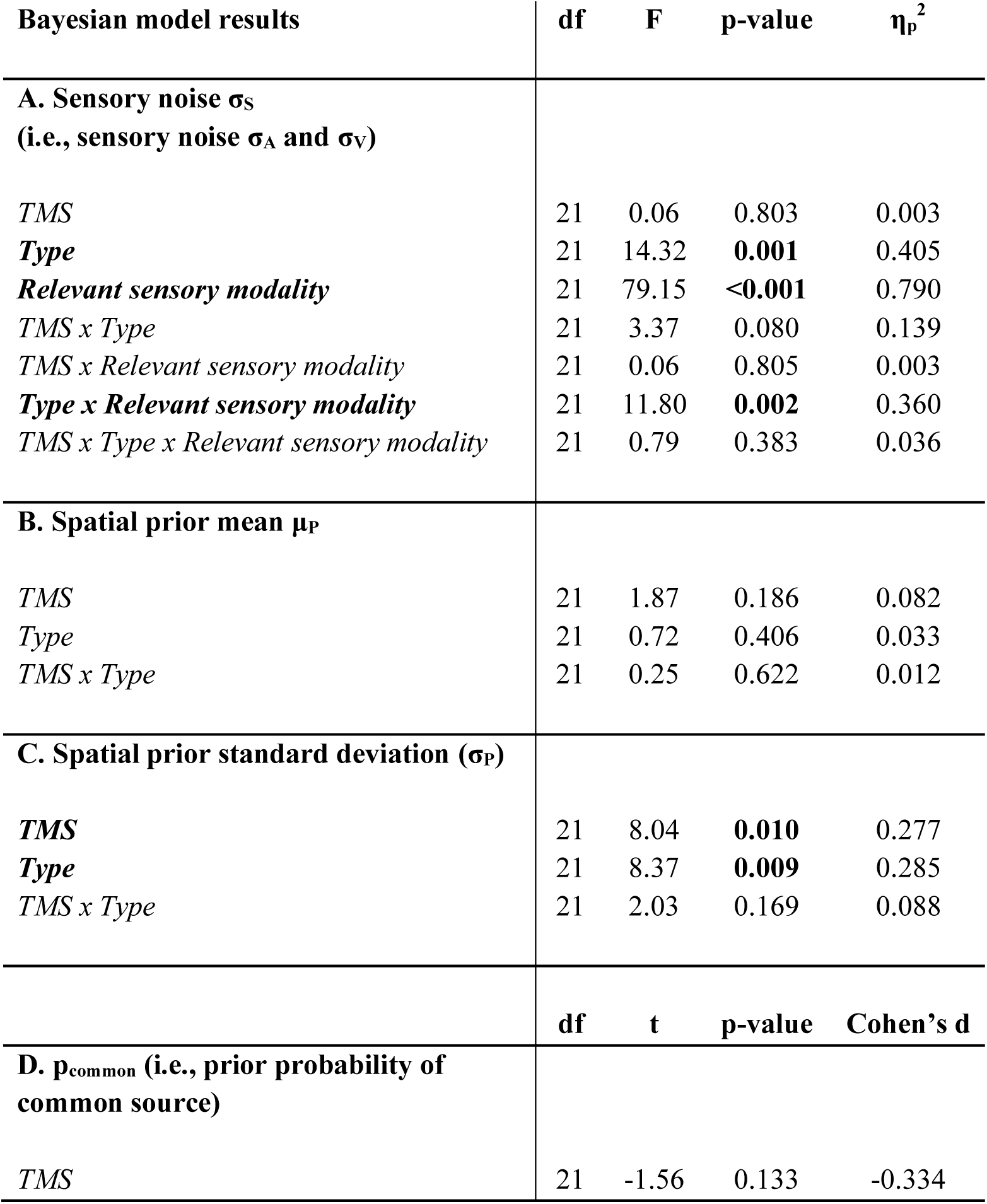
Summary of results of ‘Bayesian causal inference model’ (BCI) analyses. *TMS*: sham-TMS vs. IPS-TMS; *Relevant sensory modality*: report Auditory vs. Visual stimuli; *Type*: Auditory/Visual vs. Audiovisual congruent. Significant factors and their *p*-values are highlighted in bold.

#### IPS-TMS increases spatial prior’s standard deviation (σ_P_) but leaves the mean(μ_P_) unaffected

A 2 (*TMS*: sham-TMS vs. IPS-TMS) × 2 (*Type*: Auditory/Visual vs. Audiovisual) repeated measures ANOVA on μ_P_ revealed no significant main effects or interactions, mirroring our model-free results (**Table 3B**). Importantly, however, the same analysis on σ_P_ revealed a significant main effect of TMS (**Table 3C** and **Fig.2D**). In line with our model-free analyses, IPS-TMS significantly increased the standard deviation of the spatial prior, thereby decreasing its weight on observers’ spatial percepts. These Bayesian modelling results support our tentative conclusions from the model-free analysis, namely that IPS-TMS broadens observers’ response distribution by reducing the precisions of their spatial priors.

#### No effects of IPS-TMS on prior probability of a common source (p_common_)

A paired-sample t-test (*TMS*: sham-TMS vs. IPS-TMS) revealed no significant difference between sham-TMS and IPS-TMS on observers’ causal prior or tendency to bind audiovisual signals (**Table 3D**). Again, this null-effect converges with the model-free analysis showing no significant IPS-TMS effect on the crossmodal bias.

In summary, the findings from the model-based analysis corroborate those from the model-free analysis. IPS-TMS selectively increases the standard deviation or variance of observers’ spatial prior without affecting observers’ prior mean or causal prior or auditory/visual noise. Since we estimate observers’ spatial prior collectively from all trials across the entire experiment, it reflects observers’ central response bias - a general tendency to perceive and report locations skewed towards the overall mean of all stimulus locations in their left hemifield.

In addition to this central response tendency, extensive research has shown that observers’ perceptions are often biased towards their previous perceptual decisions. From a Bayesian perspective these serial dependencies across trials arise when observers dynamically adapt their priors in response to the changing sensory statistics (21, 27–31). Critically, observers have also been shown to adapt their priors in experiments in which the statistical structure remains constant and stimuli are presented in a randomized order (31). This raises the question of whether observers’ spatial decisions in our study are also biased towards their previously perceived stimulus location and whether these dynamic biases decrease following offline low frequency IPS-TMS.

### IPS-TMS reduced the influence of observers’ prior spatial decision on their perceived stimulus locations

We estimated the influence of observers’ previous spatial decisions on their current perceived spatial locations using a linear regression model which predicted observers’ perceived stimulus locations by the current true auditory and visual locations and their previous spatial decisions (for full model description, **Materials and Methods**). As expected, observers’ spatial decisions were biased towards their previously perceived sound and flash locations consistently across all conditions, albeit to varying degrees. A 2 (*TMS*: sham-TMS vs. IPS-TMS) × 2 (*Relevant sensory modality*: report Auditory vs. Visual stimuli) × 2 (*Type*: Auditory/Visual vs. audiovisual) repeated measures ANOVA revealed significant main effects of stimulus type and relevant sensory modality (**Table 1E** and **Fig.3B**). Critically, we observed significant interactions of TMS with both stimulus type and sensory modality. As illustrated in **Fig.3B**, IPS-TMS, compared to sham-TMS, diminishes the influence of observers’ preceding spatial decisions on their current perception of stimulus locations. This effect is particularly pronounced for trials with auditory report, probably because of the greater auditory noise. In accordance with the principles of precision weighted integration, greater sensory noise amplifies the relative influence of prior beliefs, thereby making the modulatory effects of TMS more apparent.

Collectively, our results suggest that IPS-TMS reduces the influence of observers’ prior central tendency and their previous spatial decisions on their current perception of stimulus locations.

## Discussion

A wealth of research has shown that human observers combine prior knowledge with sensory signals weighted by their precisions in a manner consistent with Bayesian inference (6, 15). However, the neural basis of priors and their uncertainties remains unclear. In this study, we employed psychophysics, Bayesian modelling, and low-frequency TMS to investigate the causal role of right anterior IPS in the integration of spatial and/or causal priors (i.e., common vs. separate causes) with auditory/visual signals (i.e., likelihoods) for spatial decisions.

We presented participants with audiovisual signals at varying spatial disparities and locations limited to their left hemifield, because right IPS-TMS is likely to have distinct effects on stimulus processing in ipsi- and contralateral hemifields (32–35). Despite these experimental modifications, our study replicated the key findings of previous research in audiovisual spatial decision making: observers located visual signals with greater precision than auditory ones. Consequently, and in accordance with precision-weighted integration, a disparate visual signal biased observers’ perceived sound location but the reverse was not observed (36–38). As shown in **Fig.3A**, the audiovisual weight index for auditory reports deviated from zero indicating stronger visual influences, while it approached one indicating purely visual influence for visual reports. Crucially, these audiovisual biases and crossmodal influences were substantially reduced for large spatial disparities. Thus, in line with Bayesian causal inference, observers integrated signals when they were likely to arise from common causes, but segregated them otherwise (6).

We next investigated whether and how IPS-TMS influences the brain’s integration of spatial priors with auditory/visual inputs. According to Bayesian principles, observers should integrate prior knowledge with sensory inputs weighted by their relative precisions (i.e., inverse of noise, or uncertainty), giving a lower weight to less reliable information. In line with the notion that perceptual inference arises along the cortical hierarchy with higher areas providing prior information on lower sensory areas, we expected parietal cortices, situated at the top of the spatial processing hierarchy, to furnish spatial priors about the spatial stimulus distribution (11). Consequently, low frequency inhibitory TMS should reduce the influence of observers’ prior central tendency, resulting in a wider response distribution. In support of this hypothesis, our model-free analysis revealed a significantly wider spatial response distribution after IPS-TMS compared to sham-TMS (**Fig.2B**), in the absence of changes in auditory or visual noise (**Fig.2A**). While this model-free analysis provides tentative evidence for an IPS-TMS effect on observers’ reliance or weight assigned to spatial priors, these results need to be followed up with a model-based analysis that can formally dissociate spatial priors and sensory evidence by fitting a Bayesian causal inference model to observers’ spatial response distributions. Mirroring the findings from the model-free analysis, Bayesian modelling confirmed that IPS-TMS increased the spatial prior’s variance (**Fig.2D**), without affecting its mean or observer’s auditory or visual noise (**Fig.2C**). Collectively, our model-free and model-based analyses thus provide convergent evidence that offline low-frequency TMS increases the variance of observer’s spatial prior. As a consequence, observers rely less on their spatial prior or central response tendency leading to a widening of their response distribution.

In addition to this central response tendency and long-term biases induced by the stimulus distribution (11, 12, 39), extensive research has shown that observers exhibit dynamic biases towards their previous perceptual decisions across a wide range of perceptual stimuli and tasks (28–31). From a Bayesian perspective, these serial dependencies across trials can be attributed to observers’ priors that dynamically adapt to changes in the sensory statistics. Importantly, this dynamic adaptation has been observed even when stimuli are drawn from a constant distribution and presented in a random order – a finding that has been explained by observers’ prior belief in a non-stationary, i.e., dynamic, world (21, 27–31). Observers in our study exhibit a similar behavior. Despite stimuli being sampled independently from the four possible locations with equal probability across trials, observers’ percepts are skewed towards their previous spatial decisions. This impression was confirmed by our regression analysis: regressors encoding observers’ previous spatial decisions were able to predict observers’ current spatial percepts above and beyond the current auditory and/or visual stimulus locations alone. These dynamic biases were observed mainly for auditory report trials that are associated with greater sensory noise (**Fig.2A**). The lower auditory spatial precision thus helps to uncover the biasing effect of spatial priors, while the high spatial precision of vision obscures effects of prior spatial beliefs. These prominent dynamic biases towards previous spatial decisions - particularly under auditory report - raise the question whether they can be reduced by offline low-frequency IPS-TMS. Indeed, we observed a statistically significant reduction of these spatial biases by IPS-TMS compared to sham-TMS, particularly for auditory report trials (**Fig.3B**).

Collectively, our results suggest that TMS applied to anterior parietal cortices selectively diminishes the influence of observers’ prior central tendencies and their previous spatial decisions on their current spatial percepts. Offline IPS-TMS might directly affect the representations of spatial priors. For instance, it may increase the variance of observers’ spatial priors, thereby decreasing their weight in perceptual inference processes and resulting in lower central response biases. Alternatively, IPS-TMS could impair the brain’s ability to dynamically adapt and encode spatial priors across trials. Previous modelling research has shown that dynamic adaptation of perceptual priors can explain both the serial dependencies of their percepts across trials as well as observers’ central response tendency (30). Therefore, we favor the latter explanation as the more parsimonious one.

Interestingly, we observed no TMS effect on the spatial prior’s mean, in contrast to its effect on the spatial prior’s variance. This result may at first sight be surprising given the well-known neglect symptoms after right parietal lesions. Patients with hemispatial neglect and right hemispheric lesions typically fail to detect stimuli in their left hemifield or exhibit lateralized spatial biases in a variety of tasks (40–44). Moreover, these behavioral changes were also observed following repetitive TMS to posterior parietal cortices (32, 34, 35, 45–50). Critically, those earlier studies used a detection task and presented visual stimuli in both hemifields. Others stimulated both visual cortices simultaneously using line bisection tasks, thereby inducing inter-hemispheric competition.

Our study, however, intentionally avoided this inter-hemispheric competition by restricting stimulus locations to the hemifield contralateral to IPS-TMS. Moreover, previous research primarily focused on visual signals (though see (51)). In contrast, our study revealed TMS effects on the dynamic influences of spatial expectations mainly for auditory signals that are associated with lower spatial precision and thereby allow us to detect even subtle effects of prior spatial beliefs.

Our results align perfectly with the extensive neuroimaging evidence suggesting that parietal cortices play a critical role in integrating audiovisual spatial inputs with prior spatial beliefs into spatial representations to guide perceptual decisions (e.g., 52–54). In particular, a recent fMRI study demonstrated that parietal cortices encode auditory and visual stimulus locations, their spatial congruency and motor output, while dorsolateral prefrontal cortices coded observers’ causal decisions, i.e., whether auditory and visual signals originate from the same or different sources (23).

In conclusion, our study demonstrates that, compared to sham-TMS, IPS-TMS reduces the influence of prior spatial expectations on observers’ current spatial perception thereby leading to a wider spatial response distribution. These findings support a causal role of anterior IPS in dynamically updating and encoding spatial, but not causal, priors. Future studies are required to assess the potential causal role of dorsolateral prefrontal cortices in encoding observers’ causal priors in audiovisual spatial perception.

## Materials and Methods

### Participants

Twenty-five healthy participants (14 females; mean age = 25.24 years; range 19-44 years) participated in the study. All participants had normal or corrected to normal vision, reported normal hearing, had no history of neurological or psychiatric illness and had no contraindications to TMS as listed in the safety questionnaire (55). All participants were right-handed, according to the Edinburgh Handedness Inventory ((56); mean laterality index, 92; range 70-100). Three subjects were excluded after the initial screening study, because their localization performance did not meet the inclusion criteria (**Design and Procedure**). Hence, twenty-two participants (13 females; mean age = 24.23 years; range 19-32 years) took part in the TMS experiment. This sample size was determined based on previous neurostimulation studies investigating multisensory processes (57–62).

All participants gave informed written consent to participate in the experiment. The study was approved by the research ethics committee of the University of Birmingham (approval number: ERN_15-1458P) and was conducted in accordance with the principles outlined in the Declaration of Helsinki.

### Stimuli and Apparatus

Auditory and visual stimuli were presented at four locations along the azimuth at -8°, -16°, -24° and -32° visual angle. The stimulus presentation was limited to the left hemifield because right parietal TMS is known to exert opposite effects for the ipsi- and contralateral hemifields (32–35). Auditory stimuli were bursts of white noise of 40 ms duration (with 5 ms onset and offset ramps) presented at approximately 70 dB SPL via four pairs of external speakers (MEMTEQ, Shenzhen, China) aligned vertically above and below the left monitor. The four pairs of speakers were located at -8°, -16°, -24° and -32° of the participants’ midline (i.e., on the left of the fixation light). Visual stimuli (“flashes”) were white discs (radius: 2.5° visual angle) of 40 ms duration presented on a black background. They were presented at -8°, -16°, -24° and -32° azimuth and 0° elevation in correspondence to the mean location of each pair of speakers (**Fig.1B**).

During the screening session and the four sessions of the TMS experiment, participants sat in front of two LCD monitors (1920 × 1080 resolution, 60 Hz refresh rate, 24” Iiyama ProLite B2481HS). The two monitors were tilted by -25° (monitor on the left) and +25° (monitor on the right) along the azimuth, thus forming an angle of 130°. This was done so that the auditory and visual stimulus locations at -8°, -16°, -24° and -32° could be considered to lie approximately on a circle. Throughout the entire experiment, a fixation light (0.8° diameter) was presented in the middle of the two monitors at 0° of vertical visual angle and at a viewing distance of approximately 50 cm (**Fig.1B**).

Stimuli were presented using Psychtoolbox version 3 ((63); www.psychtoolbox.org), running under Matlab R2017a (Mathworks Inc., Natick, MA, USA) on a Windows machine. Participants fixated throughout the entire experiment, resting their chin on a chinrest. Their fixation was monitored using the EyeLink eyetracking system (SR Research, 2007, 2000 Hz sampling rate; **Fig.1B**). Participants responded by pressing one of four keys of a keypad with four fingers (index, medium, ring and little finger) of their right hand (i.e., dominant hand). The order of the keys corresponded to the order of the stimuli location, from left to right.

### Design and Procedure

The study included 5 days. Day 1 included familiarization with the stimuli and task and an assessment of participants’ auditory and visual localization performance. Depending on whether their performance exceeded the inclusion criterion, participants would continue with the main IPS-TMS/sham-TMS experiment, from day 2 to day 5.

#### Practice and screening session (day 1)

Participants were familiarized with the mapping from stimulus location to response button. To train them on auditory and visual localization, they were presented with blocks of sounds and flashes sampled randomly from the four different locations along the azimuth. On each trial, observers reported the stimulus location and were given corrective feedback (i.e., green or red light at the fixation cross). At the end of each block, observers were shown their average performance accuracy during the previous block and received verbal feedback about their fixation performance. After this initial familiarization run, participants localized unisensory auditory and visual stimuli, randomly sampled from the four possible locations without corrective feedback (45 trials per location × 4 locations × 2 sensory modalities = 360 trials in total, presented in 3 auditory and 3 visual blocks). If participants’ auditory and visual localization performance passed our inclusion criterion (<8° for sounds and <4° for flashes, defined based on prior pilot study with 10 participants), they continued with the main TMS experiment on days 2 to 5. Localization performance was computed as the root mean square error (RMSE) between participant’s reported- and the true-physical stimulus location, averaged across locations. 22 out of the 25 participants passed the inclusion criterion. The mean group RMSE (including all 25 participants) was 5.3° ± 0.2° (across subjects mean ± SEM) for sounds and 3.2° ± 0.2° (across subjects mean ± SEM) for flashes.

#### Experimental design of main TMS experiment (days 2-5)

In a spatial ventriloquist paradigm, participants were presented with synchronous, spatially congruent or disparate visual and auditory signals. On each trial, visual and auditory locations were independently sampled from four possible locations along the azimuth (-8°, -16°, -24°, -32°) leading to four levels of audiovisual spatial disparity (i.e., 0°, 8°, 16° or 24°, **Fig.1A**). Thus, the 4 × 4 × 2 factorial design manipulated (i) the *location of the auditory stimulus* (-8°, -16°, -24°, -32°, absent), (ii) the *location of the visual stimulus* (-8°, -16°, -24°, -32°, absent), (iii) *relevant sensory modality* block (report auditory or visual signals) and (iv) *TMS* (IPS-TMS or sham-TMS). In addition, unisensory auditory and visual stimuli were presented from these four locations. In an intersensory selective attention task, observers located either the auditory or the visual signal. On each trial (total SOA: 2040 ms) observers were first presented with fixation light alone (500 ms duration), followed by a brief sound, flash, or sound-flash (stimulus duration: 40 ms), and finally the fixation light alone (1500 ms as response interval). Prior to the main experiment, we applied either sham-TMS or IPS-TMS on different days (2 days sham-TMS + 2 days IPS-TMS, in an alternated fashion). The order of sham-TMS and IPS-TMS were pseudorandomized across participants.

Stimulus location was randomized across trials. Relevant sensory modality was pseudorandomized over 12 blocks (i.e., 6 visual + 6 auditory report) of 60 trials each (block duration approximately 122 s) within each session and counterbalanced across subjects. Prior to each block, a cue (duration: 5000 ms) informed participants about the task-relevant sensory modality (**Fig.1C**). The order of TMS sessions was counterbalanced across subjects. The experiment included 60 trials × (6 visual + 6 auditory) relevant sensory modality blocks × 4 TMS sessions (2 sham and 2 IPS) = 2880 trials in total (= 18 trials × 20 conditions [16 audiovisual + 4 auditory/visual] × 2 relevant sensory modalities [auditory and visual] × 2 TMS [sham and IPS] × 2 sessions).

On each day and before the start of the main experiment, participants were briefly re-familiarized with the stimuli in three unisensory blocks with corrective feedback, followed by the spatial ventriloquist task without corrective feedback (20 trials for each relevant sensory modality condition).

### Control analysis: Eye movements

We recorded participants’ eye movements to exclude the possibility that results were confounded by eye saccades. We employed an EyeLink 1000 Plus system (SR Research, 2007) with sampling rate of 2000 Hz. Eye data were on-line parsed into saccade, fixation, and eye blink, using the EyeLink 1000 Plus software (velocity threshold = 30°/sec, acceleration threshold = 8000°/sec^2^, motion threshold = 0.15°). Separately for each condition, we computed the number of saccades averaged across trials. We recorded 2.67 saccades per trial (SEM = 0.25), averaged across participants and conditions. A 2 (*TMS*: sham-TMS vs. IPS-TMS) × 2 (*Session*: first vs. second) × 2 (*Modality*: auditory vs. visual) × 4 (*Location*: 8°, 16°, 24°, 32°) repeated measures ANOVA on mean number of saccades over unisensory auditory/visual trials revealed only a significant effect of Modality, with more saccades for auditory than visual trials (F(1,21) = 7.24, *p* = 0.014, η_p_^2^ = 0.26). A 2 (*TMS*: sham-TMS vs. IPS-TMS) × 2 (*Session*: first vs. second) × 4 (*A location*: 8°, 16°, 24°, 32°) × 4 (*V location*: 8°, 16°, 24°, 32°) repeated measures ANOVA on mean number of saccades over audiovisual trials revealed no significant effects or interactions (*p* > 0.05). These results (i.e. no TMS effect on eye movements) conform that TMS effects on observers’ priors cannot be attributed to eye movement confounds.

### MRI acquisition

A 3T TIM Trio System (Siemens, Erlangen, Germany) was used to acquire a high-resolution structural image (TR = 8.4 ms, TE = 3.8 ms, flip angle = 8°, FOV = 288 mm × 232 mm, image matrix = 288 × 232, 175 sagittal slices acquired in ascending direction, voxel size = 1 mm × 1 mm × 1 mm). Anatomical MRI data were processed with statistical parametric mapping (SPM12; Wellcome Department of Imaging Neuroscience, London; www.fil.ion.ucl.ac.uk/spm, (64)).

### TMS site and procedure

Two TMS conditions were included in the experiment: stimulation of the right anterior IPS as experimental condition and sham stimulation as control condition. The right anterior IPS was selected because brain areas IPS3 and IPS4 have previously been implicated in audiovisual spatial processing in a similar spatial ventriloquist paradigms (16).

We defined individual stimulation coordinates for each participant by (1) inverse transforming the right IPS3 and IPS4 probabilistic maps from (65) from MNI to native space using the parameters obtained from spatial normalization of individual T1-weighted structural MRI images, (2) identifying the coordinates (in native space) of the voxel with maximal probabilities summed over the inverse transformed IPS3 and IPS4 probabilistic maps (SPM12 MNI single subject’s coordinates of the summed maximal probabilities: × = 20, y = -64, z = 60) and (3) computing the stimulation entry as the intersection between the scalp and the shortest vector connecting the individual coordinates to the scalp.

We applied offline low-frequency (1 Hz) repetitive TMS in one single continuous train for 30 minutes using a MagPro X100 stimulator (MagVenture, Denmark) and a figure-of-eight coil (MCF-B70, MagVenture Company, Denmark) positioned over the stimulation site with the coil handle held posterolaterally at about 45° from the mid-sagittal axis. This offline stimulation protocol has been shown to decrease cortical excitability (66) with associated behavioral effects (32, 67). Moreover, it avoids TMS-specific auditory and somatosensory side effects on sensory cortices (68). Because low-frequency repetitive TMS (i.e., ≤1 Hz) is expected to reduce cortical excitability for about 50-100% of the stimulation duration (69, 70), experimental task duration was also set to 30 mins, i.e., matched to the TMS stimulation duration.

Following (55) the stimulation intensity was set at 100% of the individual resting motor threshold, determined as the minimum TMS intensity applied over the right motor cortex required to evoke a visible twitch of the participant’s left hand in approximately five out of ten consecutive pulses. Mean TMS intensity across participants was 33% stimulator output (range 24%-42%).

The positioning of the TMS coil and real time monitoring of coil position was performed with the help of a neuronavigation system (Localite, Bonn, Germany). To maximize stimulation accuracy, the TMS coil was attached to a mechanical arm and participants’ head movements were minimized with cushioned support around their head. Participants were provided with earplugs and were allowed to close their eyes.

Sham-TMS was performed by positioning the coil on the same spot as for IPS-TMS but tilting it to 90° with the coil wings touching the scalp. Such positioning has been shown to produce similar acoustic effects and sensations as real TMS, without inducing biological activity in the cortex (71). During sham-TMS, stimulation intensity was set to the same level as for IPS-TMS.

IPS-TMS (2 sessions over 2 days) and sham-TMS (2 sessions over 2 days) order was alternated, with at least 24 hours between IPS-TMS and sham-TMS sessions and seven days between the two IPS-TMS sessions (55). To measure potential differences between subjective effects of sham-TMS and IPS-TMS stimulation, after each session we asked participants to answer the question: *During stimulation, how often did you experience:* (*1*) *Neck pain,* (*2*) *Headache* (*3*) *Pinching sensation where the coil was positioned,* (*4*) *Heat sensation where the coil was positioned,* (*5*) *Loud clicking noise,* (*6*) *Fatigue,* (*7*) *Faint feeling,* (*8*) *Drowsiness*, by rating each item on a 10-point Likert scale (0: never, 10: the whole time). To evaluate whether participants became aware of the stimulation difference, after the final session participants completed a questionnaire where (1) they indicated whether they noticed any difference between sessions during the TMS stimulation and (2) after being informed about the application of two different types of stimulation, they were asked to indicate (or guess) in which sessions they believed sham-TMS was applied. A comparison of the Likert scale responses on subjective effects experienced after sham-TMS and IPS-TMS (2-tailed Wilcoxon Signed Ranks tests) did not show significant difference between sham-TMS and IPS-TMS in any of the questions (*p* > 0.05, not corrected for multiple comparisons). Nine out of 22 participants reported that they noticed a difference between stimulation conditions once they were made aware of it, and five of them correctly guessed the relative sessions where sham-TMS and IPS-TMS were applied. Additional analyses including IPS-TMS vs. sham-TMS awareness as a regressor of no interest replicated our initial findings and will therefore not be further discussed.

## Data analysis

### ‘Model-free’ and ‘model-based’ characterization of the response distributions

We assessed the effect of IPS-TMS on observers’ spatial localization behavior using two complementary approaches: 1. Our ‘model-free’ (MF) analyses characterized observers’ response distributions (pooled over days) in terms of the sensory noise standard deviation (σ_MF_S_), mean (μ_MF_O_), spread (σ_MF_O_) and crossmodal biases as ‘model free indices’. 2. Our ‘model-based’ (BCI) analysis fits the Bayesian Causal Inference model to participants’ response distribution to obtain spatial (μ_P_, σ_P)_ and causal prior (p_common_) along with sensory noise standard deviations (σ_A_, σ_V_ that are combined into σ_S_) as parameters (**Table 2**). At the group (i.e., between subject level), both analysis approaches entered these model-free indices or model-based parameter estimates into general linear models (e.g., repeated measures ANOVA).

#### ‘Model-free’ (MF) analyses: sensory noise, overall mean, spread, and crossmodal bias

We computed several model-free indices to characterize observers’ response distributions and their audiovisual interactions.

*Sensory noise σ*_*MFS*_. We assessed observers’ auditory and visual noise in terms of the spread of their reported locations on the unisensory auditory or visual and the congruent audiovisual trials (i.e., incongruent trials were excluded from this analysis). The spread of observers’ perceived sound or flash locations can serve as a model-free proxy corresponding to the likelihood variance in the Bayesian causal inference model. We computed the standard deviation (σ_MF_S_) of response distribution (using Bessel’s correction) separately for each of the 4 stimulus locations and the 8 conditions spanned by 2 (auditory/visual and audiovisual congruent stimuli) × 2 (report auditory vs. visual) × 2 (sham-TMS vs. IPS-TMS).

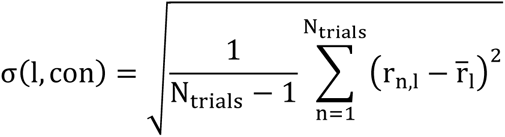

with 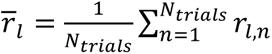, *σ*(*l*, *Con*) the standard deviation over participants’ responses, *r_n_* the response on a trial n, *N*_*trials*_ the number of trials for a particular location × condition combination.

We then pooled over the *N*_*L*_ = 4 stimulus locations to obtain a standard deviation for each condition:

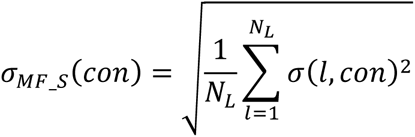

The eight *σ*_*MF*_*S*_(*con*)values (computed individually for each participant) were entered in a 2 (*TMS*: sham-TMS vs. IPS-TMS) × 2 (*Relevant sensory modality*: report Auditory vs. Visual stimuli) × 2 (*Type*: Auditory/Visual vs. Audiovisual congruent) repeated measures ANOVA.

#### Overall mean *μ*_*MF*_*O*_ and spread *σ*_*MF*_*O*_ of the response distributions

We assessed whether the mean (*μ*_*MF*_*O*_) of observers’ overall response distribution (i.e., pooled over all locations) is biased. For instance, right IPS-TMS may induce a rightward shift of observers’ perceived stimulus locations (e.g., 46, 47, 72). This response distribution’s mean can serve as a model-free proxy corresponding to the spatial prior’s mean from the Bayesian Causal Inference model.

In addition, we assessed the spread of observers’ response distribution (*σ*_*MF*_*O*_). The spread of the response distribution is influenced by both the precision (i.e., inverse of variance) of observers’ spatial prior and their sensory noise (i.e., likelihood variance). Both, a decrease in the precision of observers’ spatial prior (as a central response tendency) and an increase in the precision of their sensory noise reduces observers’ reliance on their central spatial prior-thereby increasing the spread of their response distribution. Hence, changes in the spread of observers’ response distributions need to be interpreted with caution and only together with associated changes in their sensory noise. Individually for each observer we computed the mean (*μ*_*MF*_*O*_) and spread (standard deviation; *σ*_*MF*_*O*_) of the response distributions (i.e., with trials pooled over the four stimulus locations) separately for the four conditions spanned by 2 (*TMS*: sham-TMS vs. IPS-TMS) × 2 (*Type*: Auditory/Visual vs. Audiovisual).

The spread of the response distribution was computed as standard deviation (using Bessel’s correction).

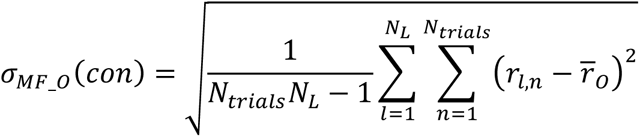

With 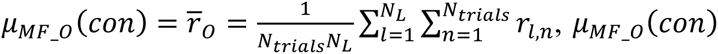 the mean, and *σ_MF_0_* (*Con*) the standard deviation over all participants’ responses *r*_*n*_ in a particular condition, *N*_*trials*_ the number of trials for a particular location (l) × condition (con).

*μ*_*MF*_*O*_(*con*) tand *σ*_*MF*_*O*_(*Con*) were entered separately in two 2 (*TMS*: sham-TMS vs. IPS-TMS) × 2 (*Type*: Auditory/Visual vs. Audiovisual) repeated measures ANOVA. To ensure that potential TMS-effects on the spread of the response distribution were not driven by crossmodal biases, we repeated this analysis constrained to the audiovisual congruent trials alone.

#### Crossmodal bias

To evaluate the effect of IPS-TMS on how audiovisual signals are integrated, we computed the audiovisual weight index W_AV_ (n.b., this can only be computed for the audiovisual incongruent trials). W_AV_ was computed as the normalized difference between the mean response location (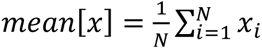) of the incongruent AV condition (Alocation ≠ Vlocation) minus the mean response location of the congruent AV condition (Alocation = Vlocation). Note that the AV congruent stimulus location matched the auditory stimulus location of the AV incongruent condition:

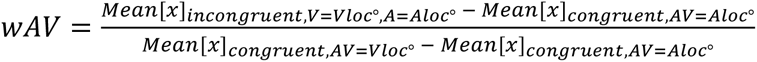

W_AV_ = 1 indicates that the observer’s spatial report relies exclusively on the location of the visual signal. W_AV_ = 0 indicates that the observer’s spatial report relies exclusively on the location of the auditory signal.

We computed W_AV_ separately for each of the 32 conditions spanned by 4 (stimulus location) × 2 (auditory/visual and audiovisual congruent stimuli) × 2 (report auditory vs. visual) × 2 (sham-TMS vs. IPS-TMS). Finally, we pooled W_AV_ over all conditions with the same absolute disparity (i.e., 8°, 16°, 24°) to obtain 12 pooled W_AV_ values for each participant: 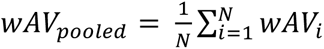 . These 12 *wAV*_*pooled*_ (for each participant) were entered into a 2 (*TMS*: sham-TMS vs. IPS-TMS) × 2 (*Relevant sensory modality*: report Auditory vs. Visual stimuli) × 3 (*Disparity*: 8°, 16°, 24°) repeated measures ANOVA.

#### Model-based analysis: Bayesian Causal Inference model (BCI)

Next, we fitted the Bayesian causal inference (BCI) model to participants’ response distributions. Briefly, the generative model of Bayesian Causal Inference assumes that common (*C* = 1) or independent (*C* = 2) causes are sampled from a binomial distribution defined by the common cause prior *p*_*common*_. For a common source, the ‘true’ location S_AV_ is drawn from the spatial prior distribution N(μ_P_, σ_P_). For two independent causes, the ‘true’ auditory (S_A_) and visual (S_V_) locations are drawn independently from this spatial prior distribution. We introduced sensory noise by drawing x_A_ and x_V_ independently from normal distributions centred on the true auditory (resp. visual) locations with variance parameters σ_A_^2^ (resp. σ_V_^2^). Thus, the generative model included the following free parameters: the common source prior p_common_, the spatial prior variance σ ^2^, spatial prior mean μ_P_, the auditory variance σ_A_^2^ and the visual variance σ_V_^2^ (5, 6, 15, 73, 74). The posterior probability of the underlying causal structure can be inferred by combining the common-source prior with the sensory evidence according to Bayes rule (6):

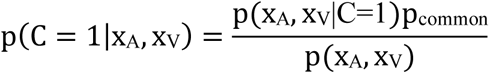

In the case of a common source (C = 1), the optimal estimate of the audiovisual location is a reliability-weighted average of the auditory and visual percepts and the spatial prior (i.e., referred to as forced fusion spatial estimate).

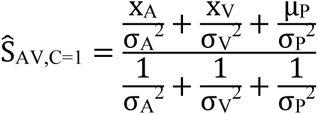

In the case of independent sources (C = 2), the auditory and visual stimulus locations (for the auditory and visual location report, respectively) are estimated independently as a reliability-weighted average of the unisensory percepts and spatial prior (i.e., referred to as unisensory auditory or visual segregation estimates):

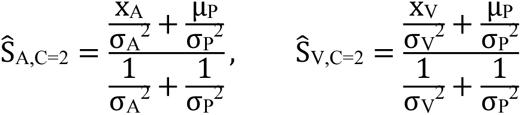

To provide a final estimate of the auditory and visual locations, the brain is thought to combine the integrated forced fusion spatial estimate with the segregated, task-relevant unisensory (i.e., either auditory or visual) spatial estimates weighted in proportion to the posterior probability of the underlying causal structures – a decision strategy referred to as model averaging (for other decision strategies, see (74)).

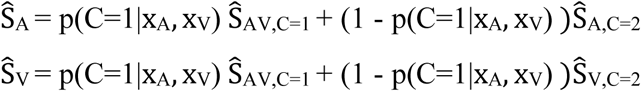

We fitted the BCI model to the audiovisual trials, separately for the sham-TMS and IPS-TMS conditions, with five free parameters: *μ_P_* and *σ_P_* (i.e., the mean and standard deviation of the spatial prior), *p_common_* (i.e., the causal prior), *σ_A_* (i.e., the auditory noise) and *σ_V_* (i.e., the visual noise) using a maximum likelihood estimation method (Table 2; BADS, (75)).

For the unisensory auditory and visual stimuli we fitted a Bayesian Causal inference model in which we fixed the p_common_ to zero, i.e., assumed full segregation. Hence, the unisensory Bayesian causal inference model included four free parameters: σ_A_, σ_V_, μ_P_, σ_P_.

Similar to our model-free analysis approach, we entered the subject-specific estimates for the spatial prior mean (μ_P_) and standard deviation (σ_P_) into separate 2 (*TMS*: sham-TMS vs. IPS-TMS) × 2 (*Type*: Auditory/Visual vs. Audiovisual) repeated measures ANOVA. Further, we entered the auditory and visual σ_A_ and σ_V_ noise parameters into a 2 (*TMS*: sham-TMS vs. IPS-TMS) × 2 (*Relevant sensory modality*: report Auditory vs. Visual stimuli) × 2 (*Type*: Auditory/Visual vs. Audiovisual) repeated measures ANOVA. Moreover, we entered the p_common_ that is only estimated for the audiovisual conditions into a paired-sample t-test (*TMS*: sham-TMS vs. IPS-TMS).

#### Serial dependencies: Influence of previous spatial percept on current spatial decisions

Extensive research has shown serial dependencies across participant’s responses throughout the experiment. Most notably, participants’ perception has been shown to be biased towards their percept of the previous trial (28–31). We examined how TMS modulates the influence of spatial percepts of the previous trial on observers’ spatial decisions on the current trial. For this, we designed regression models that included observers’ previous spatial percepts in addition to the current auditory and/or visual locations as regressors to predict observers’ spatial percepts’ on the current trials

Specifically, we estimated the influence of observers’ previous spatial percept on their current spatial percept in separate regression models for each of the 4 runs spanned by 2 (*TMS*: sham-TMS vs. IPS-TMS) × 2 (*Type*: Auditory/Visual vs. Audiovisual). The dependent variable in all those four regression models is participant’s auditory and/or visual localization response on the current trial. We computed 2 regression models for audiovisual trials and 2 regression models for unisensory trials.

The audiovisual regression model included the following predictors: the true auditory location on trials with auditory report separately for AV disparities = 0°, 8°, 16°, 24° (predictors 1-4), the true visual location on trials with visual report separately for AV disparities = 0°, 8°, 16°, 24° (predictors 5-8), the true auditory location on trials with visual report separately for AV disparities = 8°, 16°, 24° (predictors 9-11), the true visual location on trials with auditory report separately for AV disparities = 8°, 16°, 24° (predictors 12-14), the previously reported sound location on trials with auditory report (predictor 15), the previously reported visual location on trials with visual report (predictor 16), the intercept (predictor 17). Please note that because auditory and visual report trials were blocked, there were no trials with auditory report that were preceded by trials with visual report, and vice versa.

Similarly, the unisensory auditory/visual regression model included the following predictors: (1) the true auditory stimulus location on auditory trials, (2) the true visual stimulus location on visual trials, (3) the previously perceived sound location on auditory trials, (4) the previously perceived visual location on visual trials, (5) the intercept.

The subject-specific regression coefficients (i.e., parameter estimates) for the predictors encoding the response location of the previous trial (e.g., predictor 2 in the unisensory auditory model) from each of the 8 regression models were entered in a 2 (*TMS*: sham-TMS vs. IPS-TMS) × 2 (*Relevant sensory modality*: report Auditory vs. Visual stimuli) × 2 (*Type*: Auditory/Visual vs. Audiovisual) repeated measures ANOVA at the group level.

## Acknowledgments

We thank Chris Miall and Axel Thielscher for their very helpful advice on the TMS procedures. This research was funded by the ERC (ERC starting grant ‘multsens’). David Meijer is currently supported by the Austrian Science Fund (FWF, ZK-66, ‘Dynamates’).

## Author contributions

A.Z. and U.N. designed the study. A.Z. collected the data. A.Z., D.M., and U.N. analyzed the data. A.Z. wrote the first draft of the paper. A.Z. and U.N. wrote the paper. A.Z., D.M., and U.N. edited the paper. U.N. acquired the financial support for the project and supervised the project.

## References

1. C. R. Fetsch, G. C. DeAngelis, D. E. Angelaki, Bridging the gap between theories of sensory cue integration and the physiology of multisensory neurons. Nat Rev Neurosci 14, 429–442 (2013).

2. J. I. Gold, M. N. Shadlen, The neural basis of decision making. Annual Review of Neuroscience 30, 535–574 (2007).

3. W. J. Ma, J. M. Beck, P. E. Latham, A. Pouget, Bayesian inference with probabilistic population codes. Nat Neurosci 9, 1432–1438 (2006).

4. W. J. Ma, M. Jazayeri, Neural coding of uncertainty and probability. Annu. Rev. Neurosci. 37, 205– 220 (2014).

5. U. Noppeney, Perceptual inference, learning, and attention in a multisensory world. Annual Review of Neuroscience 44, 449–473 (2021).

6. K. P. Körding, et al., Causal inference in multisensory perception. PLoS ONE 2 (2007).

7. I. Vilares, J. D. Howard, H. L. Fernandes, J. A. Gottfried, K. P. Kording, Differential representations of prior and likelihood uncertainty in the human brain. Current Biology 22, 1641– 1648 (2012).

8. M. Berniker, M. Voss, K. Kording, Learning priors for Bayesian computations in the nervous system. PLoS ONE 5, e12686 (2010).

9. B. Odegaard, U. R. Beierholm, J. Carpenter, L. Shams, Prior expectation of objects in space is dependent on the direction of gaze. Cognition 182, 220–226 (2019).

10. A. A. Stocker, E. P. Simoncelli, Noise characteristics and prior expectations in human visual speed perception. Nat Neurosci 9, 578–585 (2006).

11. A. Zuanazzi, U. Noppeney, Distinct neural mechanisms of spatial attention and expectation guide perceptual inference in a multisensory world. The Journal of Neuroscience 39, 2873–18 (2019).

12. A. Zuanazzi, U. Noppeney, Modality-specific and multisensory mechanisms of spatial attention and expectation. Journal of Vision 20, 1–16 (2020).

13. F. Hong, S. Badde, M. S. Landy, Repeated exposure to either consistently spatiotemporally congruent or consistently incongruent audiovisual stimuli modulates the audiovisual common-cause prior. Sci Rep 12, 15532 (2022).

14. B. Odegaard, L. Shams, The brain’s tendency to bind audiovisual signals is stable but not general. Psychol Sci 27, 583–591 (2016).

15. L. Shams, U. R. Beierholm, Causal inference in perception. Trends in Cognitive Sciences 14, 425– 432 (2010).

16. T. Rohe, U. Noppeney, Cortical hierarchies perform Bayesian Causal Inference in multisensory perception. PLOS Biology 13, e1002073 (2015).

17. M. Aller, U. Noppeney, To integrate or not to integrate: Temporal dynamics of hierarchical Bayesian causal inference (2019).

18. Y. Cao, C. Summerfield, H. Park, B. L. Giordano, C. Kayser, Causal inference in the multisensory brain. Neuron 102, 1076–1087.e8 (2019).

19. A. Ferrari, U. Noppeney, Attention controls multisensory perception via two distinct mechanisms at different levels of the cortical hierarchy. PLoS Biol 19, e3001465 (2021).

20. G. Qi, W. Fang, S. Li, J. Li, L. Wang, Neural dynamics of causal inference in the macaque frontoparietal circuit. eLife 11, e76145 (2022).

21. T. Rohe, A. C. Ehlis, U. Noppeney, The neural dynamics of hierarchical Bayesian causal inference in multisensory perception. Nature Communications 10, 1–17 (2019).

22. T. Rohe, U. Noppeney, Reliability-weighted integration of audiovisual signals can be modulated by top-down control. Eneuro 5, 1–20 (2018).

23. A. Mihalik, U. Noppeney, Causal inference in audiovisual perception. Journal of Neuroscience 40, 6600–6612 (2020).

24. J. M. Beck, et al., Probabilistic population codes for Bayesian decision making. Neuron 60, 1142– 1152 (2008).

25. K. Friston, The free-energy principle: a unified brain theory? Nature reviews. Neuroscience 11, 127–138 (2010).

26. T. Rohe, U. Noppeney, Distinct computational principles govern multisensory integration in primary sensory and association cortices. Current Biology 26, 509–514 (2016).

27. U. Beierholm, T. Rohe, A. Ferrari, O. Stegle, U. Noppeney, Using the past to estimate sensory uncertainty. eLife 9, e54172 (2020).

28. C. W. G. Clifford, T. L. Watson, D. White, Two sources of bias explain errors in facial age estimation. R. Soc. open sci. 5, 180841 (2018).

29. J. Fischer, D. Whitney, Serial dependence in visual perception. Nat Neurosci 17, 738–743 (2014).

30. S. Glasauer, “Sequential Bayesian updating as a model for human perception” in Progress in Brain Research, (Elsevier, 2019), pp. 3–18.

31. A. J. Yu, J. D. Cohen, Sequential effects: Superstition or rational behavior? (2015).

32. C. C. Hilgetag, H. Théoret, A. Pascual-leone, Enhanced visual spatial attention ipsilateral to rTMS-induced “virtual lesions” of human parietal cortex. Nature neuroscience 4, 953–957 (2001).

33. Y. H. Kim, et al., Facilitating visuospatial attention for the contralateral hemifield by repetitive TMS on the posterior parietal cortex. Neuroscience Letters 382, 280–285 (2005).

34. G. Thut, A. Nietzel, A. Pascual-Leone, Dorsal posterior parietal rTMS affects voluntary orienting of visuospatial attention. Cerebral Cortex 15, 628–638 (2005).

35. A. Zuanazzi, L. Cattaneo, The right hemisphere is independent from the left hemisphere in allocating visuospatial attention. Neuropsychologia 102, 197–205 (2017).

36. D. Alais, D. Burr, Ventriloquist effect results from near-optimal bimodal integration. Current Biology 14, 257–262 (2004).

37. D. Meijer, S. Veselič, C. Calafiore, U. Noppeney, Integration of audiovisual spatial signals is not consistent with maximum likelihood estimation. Cortex (2019) 10.1016/j.cortex.2019.03.026.

38. T. Rohe, U. Noppeney, Sensory reliability shapes perceptual inference via two mechanisms. Journal of Vision 15, 1–38 (2015).

39. A. Zuanazzi, U. Noppeney, Additive and interactive effects of spatial attention and expectation on perceptual decisions. Scientific Reports, 1–12 (2018).

40. E. Bisiach, C. Bulgarelli, R. Sterzi, G. Vallar, Line bisection and cognitive plasticity of unilateral neglect of space. Brain and Cognition 2, 32–38 (1983).

41. M. Corbetta, et al., Neural basis and recovery of spatial attention deficits in spatial neglect. Nat Neurosci. 8, 1603–1610 (2005).

42. S. Ferber, H. O. Karnath, How to assess spatial neglect--line bisection or cancellation tasks? Journal of clinical and experimental neuropsychology 23, 599–607 (2001).

43. D. J. Mort, et al., The anatomy of visual neglect. Brain 126, 1986–1997 (2003).

44. G. Vallar, Spatial hemineglect in humans. Trends in Cognitive Sciences 2 (1998).

45. C. Babiloni, et al., Human ventral parietal cortex plays a functional role on visuospatial attention and primary consciousness. A repetitive transcranial magnetic stimulation study. Cerebral Cortex 17, 1486–1492 (2007).

46. O. Bjoertomt, A. Cowey, V. Walsh, Spatial neglect in near and far space investigated by repetitive transcranial magnetic stimulation. Brain 125, 2012–22 (2002).

47. G. Giglia, et al., Neglect-like effects induced by tDCS modulation of posterior parietal cortices in healthy subjects. Brain Stimulation 4, 294–299 (2011).

48. I. G. Meister, et al., Hemiextinction induced by transcranial magnetic stimulation over the right temporo-parietal junction. Neuroscience 142, 119–123 (2006).

49. N. G. Muggleton, et al., TMS over right posterior parietal cortex induces neglect in a scene-based frame of reference. Neuropsychologia 44, 1222–1229 (2006).

50. T. Nyffeler, et al., Neglect-like visual exploration behaviour after theta burst transcranial magnetic stimulation of the right posterior parietal cortex. European Journal of Neuroscience 27, 1809–1813 (2008).

51. F. Pavani, E. Ládavas, J. Driver, Auditory and multisensory aspects of visuospatial neglect. Trends in Cognitive Sciences 7, 407–414 (2003).

52. E. Macaluso, J. Driver, Spatial attention and crossmodal interactions between vision and touch. Neuropsychologia 39, 1304–1316 (2001).

53. S. Shomstein, S. Yantis, Control of attention shifts between vision and audition in human cortex. Journal of Neuroscience 24, 10702–10706 (2004).

54. C. Spence, “Orienting Attention: A Crossmodal Perspective” in The Oxford Handbook of Attention, (2014), pp. 1–21.

55. S. Rossi, M. Hallett, P. M. Rossini, A. Pascual-Leone, Safety, ethical considerations, and application guidelines for the use of transcranial magnetic stimulation in clinical practice and research. Clinical Neurophysiology 120, 2008–2039 (2009).

56. Oldfield, The assessment and analysis of handedness: the Edinburgh inventory. Neuropsychologia 9, 97–113 (1971).

57. C. Bertini, F. Leo, A. Avenanti, E. Làdavas, Independent mechanisms for ventriloquism and multisensory integration as revealed by theta-burst stimulation. European Journal of Neuroscience 31, 1791–1799 (2010).

58. N. Bien, S. ten Oever, R. Goebel, A. T. Sack, The sound of size. Crossmodal binding in pitch-size synesthesia: A combined TMS, EEG and psychophysics study. NeuroImage 59, 663–672 (2012).

59. N. Bolognini, C. Miniussi, S. Savazzi, E. Bricolo, A. Maravita, TMS modulation of visual and auditory processing in the posterior parietal cortex. Experimental Brain Research 195, 509–517 (2009).

60. C. D. Chambers, J. M. Payne, J. B. Mattingley, Parietal disruption impairs reflexive spatial attention within and between sensory modalities. Neuropsychologia 45, 1715–1724 (2007).

61. M. R. Kamke, H. E. Vieth, D. Cottrell, J. B. Mattingley, Parietal disruption alters audiovisual binding in the sound-induced flash illusion. NeuroImage 62, 1334–1341 (2012).

62. S. Pasalar, T. Ro, M. S. Beauchamp, TMS of posterior parietal cortex disrupts visual tactile multisensory integration. European Journal of Neuroscience 31, 1783–1790 (2010).

63. M. Kleiner, et al., What’s new in Psychtoolbox-3? Perception 36, 1–16 (2007).

64. Friston, et al., Statistical parametric maps in functional imaging : A general linear approach. Human Brain Mapping 2, 189–210 (1995).

65. L. Wang, R. E. B. Mruczek, M. J. Arcaro, S. Kastner, Probabilistic maps of visual topography in human cortex. Cerebral Cortex 25, 3911–3931 (2015).

66. F. Maeda, J. P. Keenan, J. M. Tormos, H. Topka, A. Pascual-Leone, Interindividual variability of the modulatory effects of repetitive transcranial magnetic stimulation on cortical excitability. Experimental Brain Research 133, 425–430 (2000).

67. G. S. Pell, Y. Roth, A. Zangen, Modulation of cortical excitability induced by repetitive transcranial magnetic stimulation: Influence of timing and geometrical parameters and underlying mechanisms. Progress in Neurobiology 93, 59–98 (2011).

68. F. Duecker, A. T. Sack, Pre-stimulus sham TMS facilitates target detection. PLoS ONE 8, e57765 (2013).

69. J. M. Hoogendam, G. M. J. Ramakers, V. Di Lazzaro, Physiology of repetitive transcranial magnetic stimulation of the human brain. Brain Stimulation 3, 95–118 (2010).

70. E. M. Robertson, H. Théoret, A. Pascual-Leone, Studies in cognition: the problems solved and created by transcranial magnetic stimulation. Journal of cognitive neuroscience 15, 948–960 (2003).

71. S. H. Lisanby, D. Gutman, B. Luber, C. Schroeder, H. A. Sackeim, Sham TMS: Intracerebral measurement of the induced electrical field and the induction of motor-evoked potentials. Biological Psychiatry 49, 460–463 (2001).

72. B. Fierro, F. Brighina, A. Piazza, M. Oliveri, E. Bisiach, Timing of right parietal and frontal cortex activity in visuo-spatial perception: a TMS study in normal individuals. Neuroreport 12, 2605–2607 (2001).

73. D. Meijer, U. Noppeney, “Computational models of multisensory integration” in Multisensory Perception, (Elsevier, 2020), pp. 113–133.

74. D. R. Wozny, U. R. Beierholm, L. Shams, Probability matching as a computational strategy used in perception. PLoS Computational Biology 6 (2010).

75. L. Acerbi, W. J. Ma, Practical Bayesian Optimization for Model Fitting with Bayesian Adaptive Direct Search. 1–21 (2017).

